# Rapid Development of Improved Data-dependent Acquisition Strategies

**DOI:** 10.1101/2020.09.11.293092

**Authors:** Vinny Davies, Joe Wandy, Stefan Weidt, Justin J. J. van der Hooft, Alice Miller, Rónán Daly, Simon Rogers

**Affiliations:** School of Computing Science, University of Glasgow, Glasgow, UK; Glasgow Polyomics, University of Glasgow, Glasgow, UK; Bioinformatics Group, Department of Plant Sciences, Wageningen University, 6780 PB Wageningen, The Netherlands

## Abstract

Tandem mass spectrometry (LC-MS/MS) is widely used to identify unknown ions in untargeted metabolomics. Data Dependent Acquisition (DDA) chooses which ions to fragment based upon intensity observed in MS1 survey scans and typically only fragment a small subset of the ions present. Despite this inefficiency, relatively little work has addressed the development of new DDA methods, partly due to the high overhead associated with running the many extracts necessary to optimise approaches in busy MS facilities. In this work, we firstly provide theoretical results that show how much improvement is possible over current DDA strategies. We then describe an *in silico* framework for fast and cost efficient development of new DDA acquisition strategies using a previously developed Virtual Metabolomics Mass Spectrometer (ViMMS). Additional functionality is added to ViMMS to allow methods to be used both in simulation and on real samples via an instrument application programming interface (API).

We demonstrate this framework through the development and optimisation of two new DDA methods which introduce new advanced ion prioritisation strategies. Upon application of the here developed methods to two complex metabolite mixtures, our results show that they are able to fragment more unique ions than standard DDA acquisition strategies.

## Introduction

Tandem mass spectrometry (LC-MS/MS) is increasingly used in untargeted metabolomics to aid in the annotation of unknown chemical ions. Measured fragment (MS2) spectra for unknown ions can be used to aid annotation by direct comparison against spectral databases, machine-learning assisted comparison with structural databases (e.g. SIRIUS4^1^ CFM-ID^2^) or analysis with metabolome data-mining tools such as molecular networking^3^ and MS2LDA substructure discovery. ^4^

Crucial to all of these approaches is the acquisition of MS2 data. A good MS2 acquisition strategy ought to produce spectra of a high quality for as many of the ions present in the sample as possible. There are two main approaches that are used for MS2 acquisition in metabolomics (and proteomics): Data-dependent acquisition (DDA) and Data-independent acquisition (DIA). We refer to Guo and Huan for a recent comparison and description.

DDA targets particular ions observed in MS1 survey scans for fragmentation, and is used widely in metabolomics, either in individual injections, or pooled samples. In a typical DDA scheme, the set of N ions to fragment is determined based upon the most intense ions observed in the latest MS1 survey scan. Optionally, a dynamic exclusion window (DEW) can be included that avoids fragmenting the same mass-to-charge ratio (m/z) multiple times in succession, giving lower intensity peaks more chance of fragmentation. The chosen ions are isolated and fragmented by the MS in a series of MS2 scans, which are followed by the next MS1 survey scan. The MS duty cycle therefore consists of one MS1 scan followed by up to N MS2 scans. A benefit of DDA is that the MS2 spectra emerge from the MS “ready to use” – i.e., each spectrum has been generated by fragmenting a small m/z isolation window (typically of the order of 1Da) and will therefore normally contain fragments for a single chemical species (although it is possible for multiple species to exist within a window of this size typically resulting in so-called chimeric spectra (Lawson et al., 2017)). The downsides of DDA are the limited number of ions that can be fragmented within a single injection and the somewhat stochastic nature of fragmentation (if the same injection is run twice, different ions may be fragmented).

DIA operates in a less targeted manner. Here, an MS1 scan is followed by one or more MS2 scans that do not depend on the MS1 scan. Each MS2 scan isolates a broader m/z range and can fragment many chemical species simultaneously. In theory, this means that all species in the data are fragmented although in reality it is unlikely that fragments from low intensity species will be visible in the resulting MS2 spectra. The resulting data require substantial processing to produce spectra assumed to come from a single chemical ion. This is done in software such as MSDIAL^6^ where (amongst other things) the chromatographic profile of precursor and product ions are matched. Spectra deconvolved in this way can then be used in the same manner as those produced by DDA.

There is no overall consensus as to which of these two schemes is best, and, where comparisons have been done, no clear conclusion is possible. ^5^ Although the development of improved computational tools for spectral deconvolution has allowed more applications of DIA, DDA remains a popular choice due to the high spectral quality and the fact that little or no processing is required before the spectra can be used.

Given its popularity, surprisingly little work has been done to improve DDA performance for single injections in metabolomics. Some work has looked into DDA for multiple samples, specifically DsDA^7^ for multiple injections of different samples and AcquireX^8^ for repeated injections of the same sample, but these are not useful for single injection DDA. Here, we address the problem of improving DDA coverage for a single injection, as a way of demonstrating how we can rapidly develop more general methods *in silico*.

One of the main criticisms of the performance of DDA (with respect to DIA) is its lower coverage: the proportion of ions that are fragmented. We start by theoretically computing the optimal performance for any particular injection, taking into account the uneven elution distribution of the ions. The results demonstrate that there is considerable room for improvement, motivating the development of better DDA strategies. Secondly, we describe how new strategies can be prototyped, implemented, optimised and validated using a Virtual Metabolomics Mass Spectrometer (ViMMS), ^9^ reducing the traditional need for a large amount of costly machine time. Recent novel additions to ViMMS mean that the exact same acquisition controllers can be used for both in simulation and on real hardware. Finally, we describe two new DDA acquisition strategies prototyped in this way and demonstrate, through validation on two complex samples, their improvement over traditional DDA approaches.

## Methods

### Computing Optimal DDA Performance

Computing the theoretical optimal DDA performance allows us to place an upper bound on the maximum number of fragmentation events that could occur, i.e., how many of the chemical ions present could a DDA method fragment at least once. This is not straightforward to compute as the limiting factor is often the co-elution of too many ions in certain regions of the chromatogram.

To compute optimal performance, we start by defining the ‘true’ set of chemical ions as the set of peaks picked from an .mzML file by a commonly used peak picking algorithm, such as those provided in MZmine2^10^ or XCMS.^11^ Picked peaks are represented by their bounding boxes (min and max retention time (RT) and m/z values). An MS scan schedule is created using the mean MS1 and MS2 scan times extracted from an mzML file and a fixed value of N (the number of MS2 scans for each MS1 survey scan). This results in a list of scans, and their respective scan start times. We create a bipartite graph where the two sets of nodes correspond to MS2 scans and peak bounding boxes from MS1, respectively. An edge, representing a potential fragmentation event, is added between an MS2 scan and a bounding box if the MS2 scan time is within the RT limits of the bounding box and the MS1 scan preceding the MS2 scan also has RT within the bounding box and the peak’s intensity in this MS1 scan exceeds the minimum MS1 intensity for fragmentation.

Mirroring the standard acquisition process, we compute the optimal schedule by calculating a maximum matching for this graph (see Supplementary Information S4 for more details). A matching is a subset of edges within which no two edges share an endpoint. A maximal matching is a matching such that there is no matching with more edges. A maximum matching therefore gives us the largest set of edges between scans and bounding boxes such that each scan and box is the end point for one edge. This is the largest number of peak boxes that can be targeted by the available scans. This globally optimal schedule provides useful context within which to evaluate new DDA acquisition schemes.

### Sample Preparation, Chromatography and MS Scan Settings

#### Sample Preparation

Two samples were used for our experiments to validate the performance of novel fragmentation strategies. Serum extract (QCA) was prepared from metabolite extraction of Fetal Bovine Serum (South America origin (Gibco)) by dilution 1/20 with water and addition of chloroform and methanol to the ratio of 1:1:3 (v/v/v). A beer sample (QCB) of Black Sheep Ale, 4.4%, was obtained. Sample extraction was performed by the addition of chloroform and methanol with ratio 1:1:3 (v/v/v). A vortex mixer was used to mix the extracted solution. Centrifugation was performed to remove protein and other precipitates. Finally the supernatant was removed, and the aliquot was stored at −80°C.

#### Liquid Chromatography

Chromatographic separation with HILIC was performed for all samples using a Thermo Scientific UltiMate 3000 RSLC liquid chromatography system. A SeQuant ZIC-pHILIC column was used for a gradient elution with (A) 20 mM ammonium carbonate and (B) acetonitrile. We injected 10 L of each sample into the column with initial conditions of 80% (B), maintaining a linear gradient from 80% to 20% (B) over 15 min, and finally a wash of 5% (B) for 2 min, before re-equilibration at 80% (B) for 9 min. This used a constant flow rate of 300 *μ*L/min and a constant column oven temperature of 40°C.

#### Mass Spectrometry

A Thermo Orbitrap Fusion tribrid-series mass spectrometer was used to generate mass spectra data. Full scan spectra were acquired in positive mode at a fixed resolution of 120,000 and a mass range of 70-1000 m/z. Fragmentation spectra were acquired using the orbitrap mass analyser at resolution 7,500. Precursor ions were isolated using 0.7 m/z width and fragmented with fixed HCD collision energy of 25%.

### *In silico* DDA Strategy Prototyping and Optimisation

#### Developing DDA Fragmentation Strategies

In our previous work, we introduced ViMMS,^9^ a simulator that could be used to evaluate different fragmentation strategies *in silico*. Fragmentation strategies are implemented as controllers in ViMMS. During simulation, controllers react to incoming scans and determine the next actions to perform by sending commands to the MS. Using the Top-N controller as an example, the possible acquisition commands would be whether to perform a survey (MS1) scan or to generate fragmentation (MS2) scans.

Here we have extended ViMMS by creating bridging code that allows controllers developed in ViMMS to be used directly on an actual MS. This bridge takes the form of a vendor-specific MS class in Python. Due to instrument availability, we currently support the Thermo Scientific Orbitrap Tribrid instruments through its Instrument Application Programming Interface (API) (IAPI),^12^ however the flexible design of our framework does not preclude supporting other vendors who offer real-time instrument control through an API.

Developing new methods in simulation allows us to optimise them without having to rely upon costly MS time. We therefore propose the novel controller prototyping, optimising and validating pipeline shown in Figure 1. Fullscan (mzML) data is used to seed the virtual MS.^9^ The fragmentation controller under development is implemented in the ViMMS framework in the Python programming language. It runs in the simulated environment using the virtual MS. The performance of the controller is evaluated and the best performing parameters returned. For validation on the actual instrument, the optimised parameters from the simulation are used. The results from this validation experiment are reported as the final evaluation results. The same controller code (yellow boxes in Figure 1) works with both the simulated and the actual MS.

**Figure 1:**
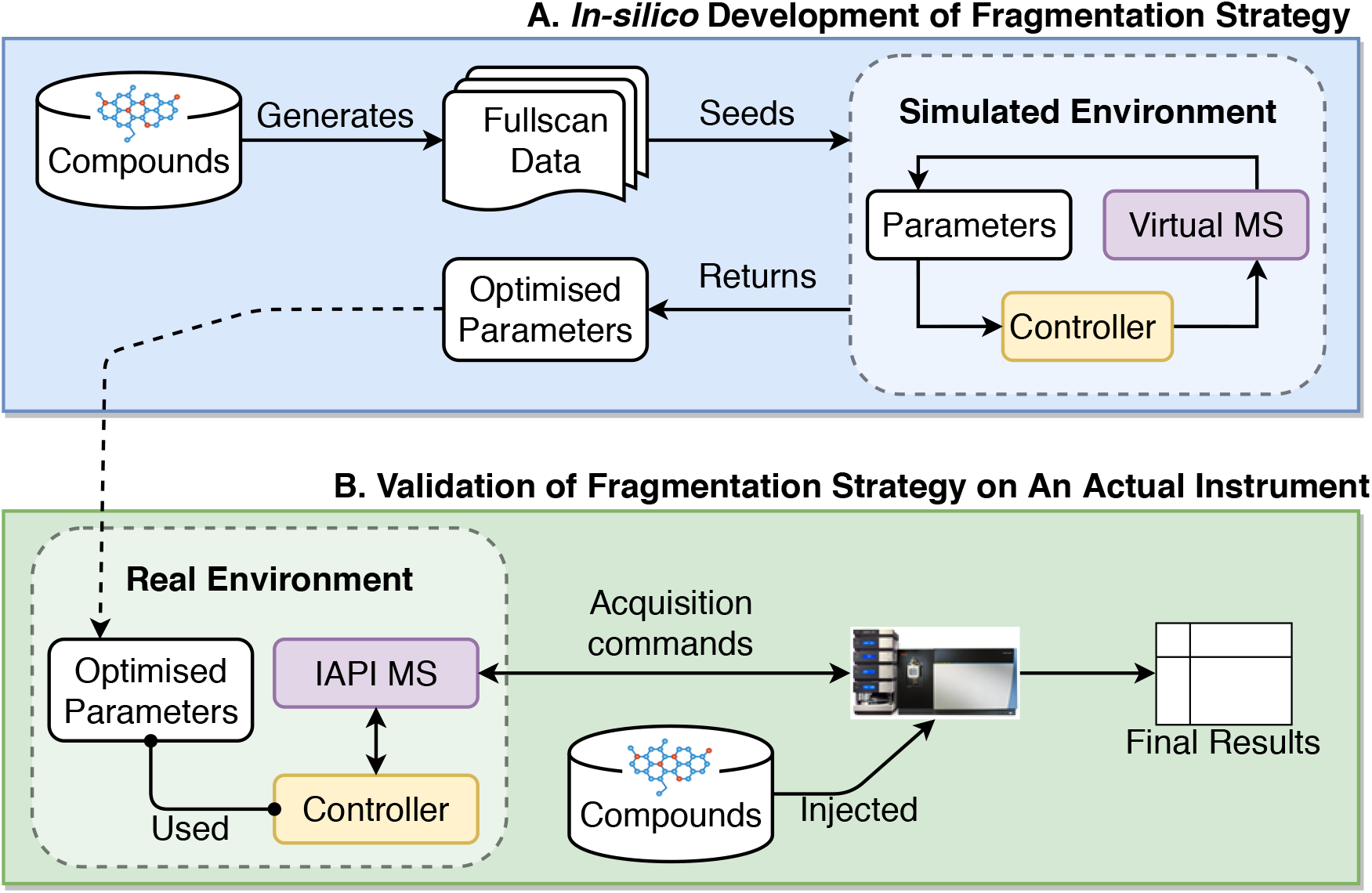
Flow diagram demonstrating the process of developing and optimising a new fragmentation strategy. **(A)** Developing, testing and optimising the fragmentation strategy *in silico*. **(B)** Validating the developed fragmentation strategy using the simulated optimal parameters on the actual instrument.

#### Performance Evaluation

We define two measures of performance to evaluate the effectiveness of different fragmentation strategies:

- *Coverage* is the number of picked peaks that contain a fragmentation event. In the absence of ground truth we use peaks picked from full-scan data acquisition.
- *Efficiency* is defined as the ratio of the number of picked peaks that are fragmented to the number of MS2 scans, i.e. how many picked peaks are, on average, targeted by one MS2 scan. A perfect value of 1.0 indicates that each fragmentation event targets one unique picked peak.

To pick peaks we use mzMine2,^10^ with parameters provided in Supplementary Table SI-1. Peaks are exported in the form of bounding boxes (m/z and RT min and max). To ensure that the results are not biased to one peak picking algorithm we also evaluated the methods using XCMS 3.6.1^11^ and Peakonly^13^ (see Supplementary Information S3). MS2 fragmentation events are checked to see which peak bounding boxes they fall into (if any). The RT range of the bounding box is defined by the first and last MS1 scans that comprise the chromatographic peak. If a fragmentation event is triggered after the last occurrence of the precursor, the resulting MS2 scan could fall outside the box, but we consider this as a hit for computing coverage. Discounting these instances made an insignificant difference to coverage.

#### Validation on Actual Instrument

For each of serum (QCA) and beer (QCB) extracts, we ran six injections: one fullscan (for evaluating coverage and efficiency), one TopN (using the controller optimised in Wandy et al.), and four injections for the new fragmentation strategies.

To compute coverage and efficiency, peaks were picked from the mzML files for the fullscan data of the two samples. With our current implementation, running methods through the Fusion IAPI requires manually starting MS acquisition once the chromatography has begun. This could lead to small time shifts between the fullscan .mzML and the .mzML from the different controllers. To ensure a fair comparison, the best results for each of the controllers are reported after performing a constant correction to the shift in retention time between −20 and 20 seconds (most shifts are far lower than this). We adopted this approach as it was the simplest way in which to align the data (any alignment method would have introduced more parameters into the analysis) and because we expected any delay in initialising to cause a small constant offset to all peaks.

### SmartROI: a Flexible Fragmentation Strategy that Targets Regions of Interest in Real-time

#### SmartROI

Our first proposed new controller is motivated by the observation that a large number of MS2 scans in the TopN controller targeted ions that were not subsequently picked as peaks. The SmartROI controller keeps track of regions of interest (ROIs) in real time and only fragments peaks within ROIs. Creation of ROIs is the first step in many peak picking methods and therefore fragmentation events outside ROIs are almost certainly wasted, see Tautenhahn et al. for an example of how to create ROIs.

SmartROI can be considered a variant on a TopN strategy in which the object being prioritised for fragmentation is the region of interest (ROI) instead of individual detected ions. As MS1 survey scans appear from the MS, the set of ROIs is updated according to the algorithm given in Tautenhahn et al.. ROIs that are not extended by the data from the MS1 scan are considered inactive and discarded. The remaining *active* ROIs are prioritised based upon intensity but only if they are available for fragmentation, determination of which is based on the following rules:

1. They must have intensity in the most recent survey scan of greater than or equal to the minimum intensity for fragmentation.
2. If they have not been fragmented before, they are available.
3. If they have been fragmented before then they are available if either of the following conditions are met:

a. Their intensity is higher by a factor *α* than when it was previously fragmented.
b. Their intensity has dropped by a factor *β* from its highest value since it was last fragmented.

Any ROI that does not meet these conditions is not available for fragmentation and will be ignored.

This strategy can be seen in Figure 2. The upper plot shows a chromatogram (x-axis retention time, y-axis intensity) and a possible set of fragmentation events using a standard TopN strategy. The dashed grey line shows the minimum intensity for fragmentation. Note that in reality, fragmentation events would depend upon the other ions eluting at the same retention time, but it is easier to understand the approaches when considered in isolation. When the intensity falls below the minimum intensity, fragmentation ceases, starting again when it rises above the threshold. In the lower plot of Figure 2, the same chromatogram is shown for SmartROI. The first fragmentation event mirrors that in the TopN. The second is slightly earlier, being triggered when the intensity has increased by *α*%. This behaviour is to ensure that we only fragment an ROI again if it has substantially increased in intensity.

**Figure 2:**
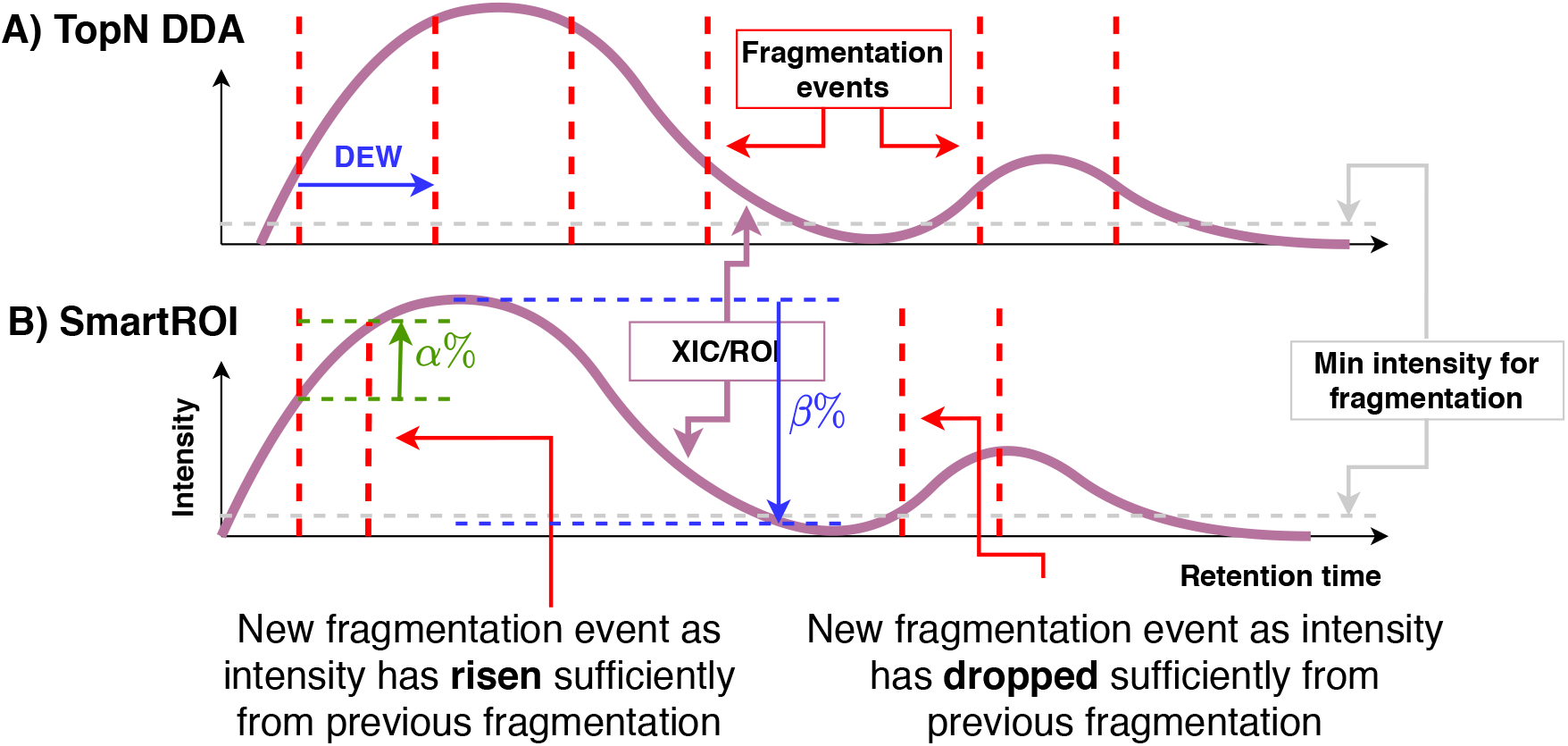
SmartROI compared with a Top-N strategy. Keeping track of an ROI in real-time allows for better targeting of MS2 events.

The SmartROI scheme then cannot fragment until the intensity has dropped by *β*% from the highest point since the previous fragmentation. However, the intensity is below the minimum intensity and so fragmentation does not occur until it has risen. The purpose of the *β*% drop is to ensure that we do not miss multiple peaks within the same ROI. The final fragmentation in the SmartROI example is triggered because the intensity has risen again by *α*%. SmartROI typically results in fewer, more precisely targeted fragmentation events than TopN.

#### Shifted SmartROI

As shown in the results (Supplementary Table SI-6), SmartROI requires additional computation when scheduling MS2 scans (updating the ROIs). We propose overcoming this by a slight variant to the controller. Rather than scheduling the next MS1 scan after N MS2 scans, we schedule it after N-1 or N-2 MS2 scans, with the remaining 1 or 2 following. This way, whilst the controller is scheduling, the MS will be acquiring MS2 scans rather than sitting idle.

### WeightedDEW: a Fragmentation Strategy with Weighted Dynamic Exclusion Scheme

WeightedDEW generalises the concept of the dynamic exclusion window. It is motivated by the problem of setting DEW width in standard TopN approaches: (i) too narrow and we waste MS2 scans repeatedly fragmenting the same ions, and (ii) too wide and we miss closely eluting peaks with similar m/z.

TopN DDA uses the intensity of the ion in the survey scan for fragmentation prioritisation. When using a DEW, peaks are excluded from repeated fragmentation as long as their m/z and RT values are still within the dynamic exclusion window of previously fragmented ions. In a standard Top-N DDA scheme, this can be thought of as prioritising ions based upon the intensity multiplied by a binary indicator (which is 0 if the ion is still excluded by DEW and 1 otherwise). The result of multiplying the precursor ion intensities and the DEW indicator terms are then used to select the TopN ranked ions to fragment. WeightedDEW generalises the binary DEW indicators to non-binary weights. It is defined by two parameters – *t*_0_ and *t*_1_. The weight for a particular ion, *w*, observed at time *t* is given by

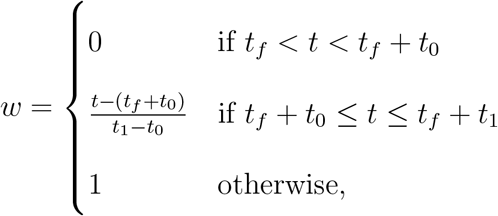

where *t*_*f*_ is the most recent time at which this m/z was fragmented. This function, for different values of *t*_1_, is shown in Figure 3a. A standard exclusion is applied for the first *t*_0_ seconds after fragmentation, after which the weight increases linearly from 0 at *t*_0_ to 1 at *t*_1_.

**Figure 3:**
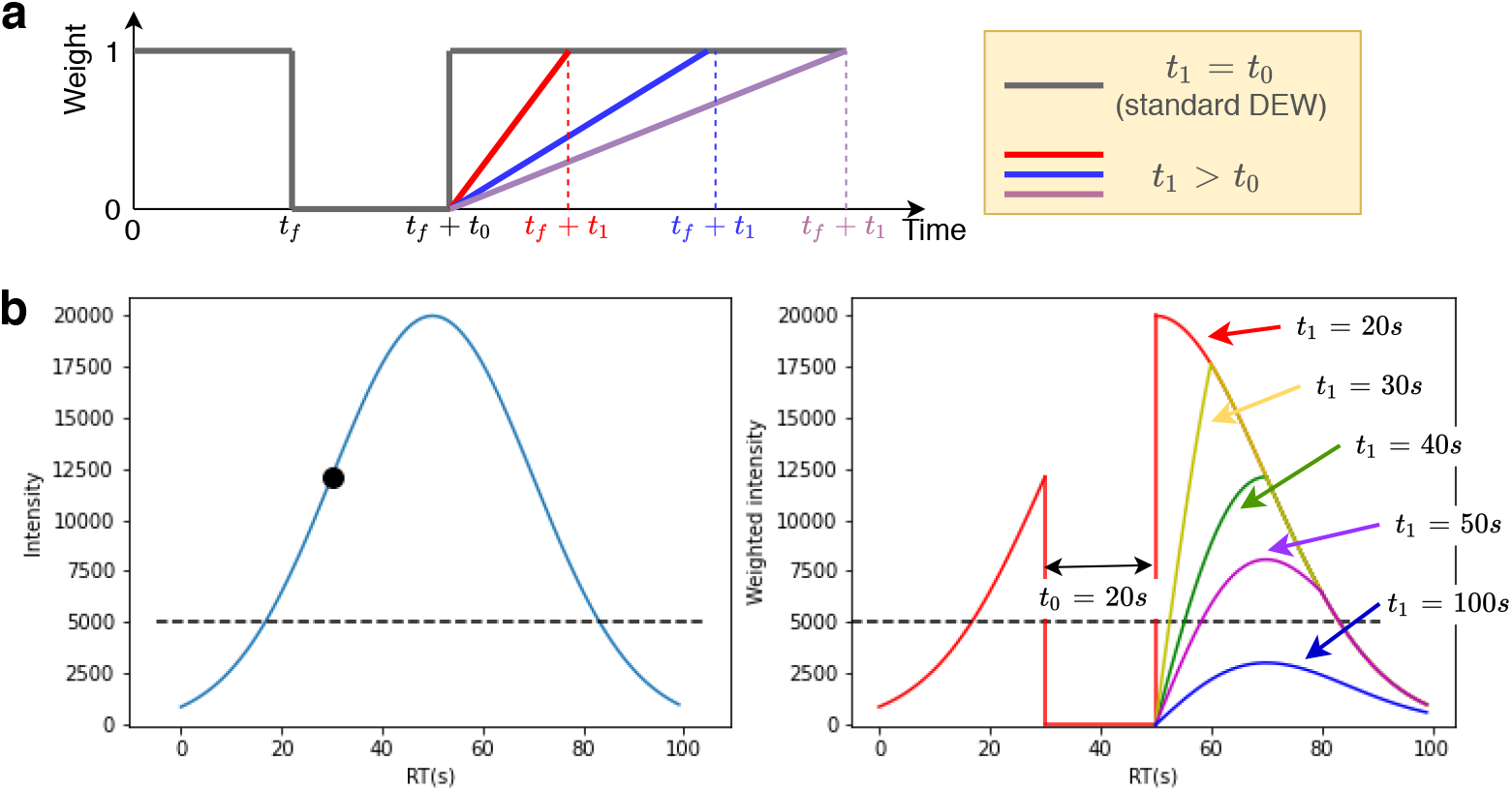
**(a)** The weight function in WeightedDEW. In standard DEW (*t*_1_ = *t*_0_) the weight is zero from the fragmentation event until *t*_0_ seconds has elapsed. In WeightedDEW, as *t*_1_ increases, the weight takes longer to return to 1. **(b)** An example chromatogram (left) showing a fragmentation event (black circle, 30s) and minimum fragmentation intensity (dashed line). The weighted intensity (right) is zero until *t*_0_ (20 seconds) has elapsed. Different curves show the effect on the weighted intensity of increasing *t*_1_.

An example chromatogram and weighted intensity can be seen in Figure 3b, with a fragmentation at 30s, *t*_0_ = 20*s* and *t*_1_ increasing from 20s to 100s. WeightedDEW down-weights chromatograms for a period after their initial exclusion. Our hypothesis is that by allowing for dynamic ‘exclusion’ to be weighted linearly as a function of time and precursor ion intensity (rather than in a binary DEW manner), the system would be able to better prioritise smaller peaks that have not yet been fragmented.

## Results

### Optimal Results

The results of our optimal analysis show that for both complex mixtures, the observed coverage from TopN DDA acquisition strategies are far from optimal, motivating the development of new methods. Optimal results were computed by picking peaks (see Supplementary Information S4) from data acquired for the serum and beer extracts in fullscan mode using scan timings taken from our own TopN method (as presented and optimised in Wandy et al.). Full results are shown in Table SI-5. In summary, for both the serum and beer extracts, the coverage of the TopN method is significantly below the optimal: 1184 (observed) v 1767 (optimal peaks) and 1509 v 2736 for serum and beer respectively. Although we would never expect to be able to reach the optimum in practice (it requires global knowledge of the peaks and when they elute), the results demonstrate the considerable room for improvement available in DDA controller design.

### Controller optimisation

Both SmartROI and WeightedDEW were optimised using a grid search for coverage in simulation (more details in Supplementary Information S7). Supplementary Figure SI-4 shows heatmaps of coverage for the serum and beer extracts for the SmartROI and Weighted DEW methods. For SmartROI, the parameter combinations *α* = 1000 and *β* = 0.1 performed well for both datasets and were chosen. For WeightedDEW, *t*_0_ = 15*s* and *t*_1_ = 120*s* were chosen. The grid search required 30 (SmartROI) and 36 (WeightedDEW) virtual injections for each of the serum and beer extracts, with each sample taking about 1 hour to produce in total – a significant time saving over running them on real equipment, demonstrating a clear advantage of optimising *in silico*.

### Validation on Instrument

After parameter optimisation, the controllers were validated on the real MS. We initially investigate the scanning frequency of the controllers. Full timing information is given in Table SI-6. Timings were computed as the difference between the scan start time in successive scans from the .mzML file in order to include the time taken to process an MS1 scan and prioritise the MS2. As expected, the SmartROI system is the slowest, with an MS1 scan taking 0.7 seconds in the beer results (compared with 0.43s for fullscan, 0.59s for TopN and 0.62s for WeightedDEW) due to the operations required to track ROIs in real time. For both serum and beer extracts, the additional processing time is the equivalent of roughly one MS2 scan motivating the development and evaluation of the shifted SmartROI controller, with shifts of 1 and 2 scans. WeightedDEW took slightly longer per MS1 scan than standard TopN. This is due to the fact that whilst TopN can greedily move from the most intense MS1 peak down until it has scheduled N MS2 scans (or runs out of non-excluded peaks), WeightedDEW has to compute the weights for all MS1 peaks above the minimum intensity threshold to ensure that it takes the topN weighted intensities into consideration. The time increase between TopN and WeightedDEW was not large enough to justify the use of a shifted controller for WeightedDEW.

Table 1 shows the performance in terms of coverage for the five controllers as well as the optimal performance as shown previously. In addition, we computed coverage based on peaks picked using XCMS and peakonly, both of which gave the same overall trends in performance with the new controllers outperforming the TopN controller (see Supplementary Information S3). In both the serum and beer extracts, the best performing controller is the WeightedDEW. SmartROI performs best with a shift of 2, as it compensates for the extra processing time required. TopN is the worst performing method in both cases. The TopN comparison used above was our own TopN controller and not the vendor TopN controller. This was due to the difficulty in comparing with the vendor controller due to the parallelisation it employs. However, for context, we compared our new fragmentation strategies against a vendor TopN controller (with identical scan parameters) and our new controllers achieved higher coverage. A more detailed description of this comparison can be found in Supplementary Information S2.

**Table 1:** Coverage (number of picked peaks fragmented) for each controller for both the serum and beer extracts, where peaks have been picked using MZmine2. The first five rows show the performance of our (Python) controllers. Numbers in brackets show the shift (in seconds) that gave this optimal performance. The final two rows give the performance of the vendor method and the optimal, computed using the bipartite graph matching, and scan timings taken from our TopN method.

| Method | Beer (6267 peaks) | Serum (4481 peaks) |
| --- | --- | --- |
| TopN | 1509 (7) | 1184 (2) |
| WeightedDEW | **2180** (10) | **1534** (2) |
| SmartROI | 1982 (4) | 1281 (-2) |
| SmartROI (shift = 1) | 2047 (4) | 1309 (1) |
| SmartROI (shift = 2) | 2106 (3) | 1318 (1) |
| Optimal (using TopN scan timings) | 2736 | 1767 |

We next consider the number of MS1 and MS2 scans produced by each method and the acquisition efficiency, shown in Table 2. We see a very wide range in the number of scans between the methods, explained predominantly by the variation in the number of MS2 scans. For the beer extract, where TopN and WeightedDEW typically create around 6000 MS2 scans, the SmartROI controllers produce far fewer, resulting in a much higher efficiency. This is explained by the relative reluctance of the SmartROI controllers to refragment the same m/z values, even after a long time has elapsed. This increased efficiency allows more MS1 scans to be produced, which is useful if these files are also being used for peak picking and relative quantification. We also hypothesise that more efficient controllers (e.g. SmartROI) would perform even better on yet more complex mixtures, where there would be more co-elution of metabolites and hence more peaks to fragment at the same time.

**Table 2:** Total number of scans, number of MS1 and MS2 scans and MS2 efficiency (the number of picked peaks that are fragmented divided by the number of MS2 scans).

| Method | Beer (6267 peaks) |  |  |  | Serum (4481 peaks) |  |  |  |
| --- | --- | --- | --- | --- | --- | --- | --- | --- |
|  | Total | MS1 | MS2 | Efficiency | Total | MS1 | MS2 | Efficiency |
| TopN | **6786** | 652 | **6134** | 0.25 | **6570** | 667 | **5903** | 0.11 |
| WeightedDEW | 6726 | 638 | 6088 | 0.36 | 6466 | 682 | 5784 | 0.27 |
| SmartROI | 4804 | 1302 | 3502 | **0.57** | 4123 | 1454 | 2669 | 0.48 |
| SmartROI (shift = 1) | 4982 | 1325 | 3657 | 0.56 | 4167 | **1505** | 2662 | 0.49 |
| SmartROI (shift = 2) | 5054 | **1343** | 3711 | **0.57** | 4212 | 1503 | 2709 | **0.50** |

## Discussion and Conclusions

Increasing MS2 acquisition coverage improves the ability to annotate ions in an LC-MS/MS analysis. However developing new acquisition methods has typically required extensive experimentation on the MS apparatus, which could be expensive and time-consuming. Here we demonstrated how new acquisition strategies can be rapidly developed and prototyped *in silico* and then validated on the machine. Additionally we introduce a framework to support this development process by extending the capability of ViMMS ^9^ so it could easily run fragmentation strategies implemented as controllers in the simulator on the real MS equipment with minimal change to the code.

Using this iterative design, prototype and validation process, we presented two new DDA acquisition strategies that both considerably outperform a conventional TopN strategy that prioritises ions for fragmentation based on intensity alone. In the first, SmartROI, we use a ROI detection algorithm commonly used for peak picking to only fragment molecules in real-time that are within ROIs and are therefore likely to be picked as peaks. In the second, WeightedDEW, we generalise the dynamic exclusion window approach to a real-valued weighting scheme allowing previously fragmented ions to smoothly rise up the priority list as their intensity remains high. In both cases, improved performance *in silico* was mapped to improved performance in reality, instilling confidence in the simulation procedures. Although the WeightedDEW controller outperformed the SmartROI in our chosen performance measure, we believe that both have utility. WeightedDEW is computationally straightforward, as demonstrated by its similar processing time to TopN, and it produces higher coverage compared to the alternatives here investigated. SmartROI requires more computational time but it also offers more direct control in how often an ROI will be fragmented. The tracking of ROIs in real time also offers the advantage of further method development. For example, it should be possible to predict, in real time, if an ROI contains a peak or not, and only fragment those predicted peaks. The increased efficiency of SmartROI also suggests that it would perform better in more crowded mixtures than those presented here. For example, background signals where the intensity values do not change much could potentially be fragmented multiple times in a standard Top-N DDA scheme, but in SmartROI it will only be fragmented once. This is possible in SmartROI even without having a prior knowledge of what the background ion is, rather it is accomplished through tracking of regions of interests in real time.

When optimising our controllers we have chosen to maximise fragmentation coverage. MS2 scan parameters have remained constant throughout, so it is not the case that we have increased coverage at the expense of data quality as would be the case if for example, reduced scan resolutions were used. All of our MS2 scans were performed in the orbitrap mass analyser to obtain high resolution fragmentation data. It would be possible to improve coverage of all methods by performing MS2 analysis in the linear ion trap mass analyser and fully make use of the possible parallelisation described in Senko et al.. The optimisation procedure proposed here is independent of any particular figure of merit: any other measure of MS2 acquisition quality could be used in place of coverage if considered more appropriate.

In addition, we have also shown how an approximate optimal limit of DDA acquisition performance for a particular mixture can be computed via a bipartite graph matching scheme. This limit provides context for acquisition analysis results: for the two complex samples analysed here, we are far from reaching these theoretical maxima, suggesting that much more optimisation is possible. At the same time, this provides a framework for future DDA and DIA method optimisation studies to perform benchmarking when applied to the samples used in their studies.

For validation on actual instruments, our proposed framework at the moment is limited to supporting the Thermo Fusion Tribrid instrument through the manufacturer’s provided IAPI. The modular nature of our software means that all controllers communicate with the instrument through bridging code and therefore the same controller implementations could easily run on different hardware if a real-time API is available from the manufacturers. For instance, Waters instruments could be supported by developing an appropriate bridge from our framework to communicate with the Waters Research Enabled Software (WREnS) API.

We conclude that there is much further improvement possible in the development of DDA acquisition strategies. We show how the use of a simulation system to optimise such strategies can rapidly lead to improvements. We demonstrate two such acquisition strategies, both exceeding performance over a TopN controller in terms of coverage (number of unique picked peaks that are fragmented).

## Acknowledgement

Vinny Davies, Joe Wandy, Stefan Weidt, Rónán Daly and Simon Rogers acknowledge EPSRC project EP/R018634/1 on ‘Closed-loop data science for complex, computationally and data-intensive analytics’. Justin J.J. van der Hooft. was partly funded by an ASDI eScience grant, ASDI.2017.030, from the Netherlands eScience Center—NLeSC.

## Supporting Information Available

The following additional information are available in the Supplementary document.

**S1** Peak picking parameters.

**S2** Scan timing analysis and comparison with vendor method.

**S3** Results for alternative peak picking methods.

**S4** Computing theoretical fragmentation bounds.

**S5** Optimality results.

**S6** Scan timings for different controller methods.

**S7***In-silico* optimisation of controllers.

## References

(1) Dührkop, K.; Fleischauer, M.; Ludwig, M.; Aksenov, A. A.; Melnik, A. V.; Meusel, M.; Dorrestein, P. C.; Rousu, J.; Böcker, S. SIRIUS 4: a rapid tool for turning tandem mass spectra into metabolite structure information. Nat. Methods 2019, 16, 299–302.

(2) Djoumbou-Feunang, Y.; Pon, A.; Karu, N.; Zheng, J.; Li, C.; Arndt, D.; Gautam, M.; Allen, F.; Wishart, D. S. CFM-ID 3.0: Significantly Improved ESI-MS/MS Prediction and Compound Identification. Metabolites 2019, 9.

(3) Wang, M. et al. Sharing and community curation of mass spectrometry data with Global Natural Products Social Molecular Networking. Nat. Biotechnol. 2016, 34, 828–837.

(4) van Der Hooft, J. J. J.; Wandy, J.; Barrett, M. P.; Burgess, K. E. V.; Rogers, S. Topic modeling for untargeted substructure exploration in metabolomics. Proceedings of the National Academy of Sciences 2016, 113, 13738–13743.

(5) Guo, J.; Huan, T. Comparison of Full-Scan, Data-Dependent, and Data-Independent Acquisition Modes in Liquid Chromatography-Mass Spectrometry Based Untargeted Metabolomics. Anal. Chem. 2020,

(6) Tsugawa, H.; Ikeda, K.; Takahashi, M.; Satoh, A.; Mori, Y.; Uchino, H.; Okahashi, N.; Yamada, Y.; Tada, I.; Bonini, P.; Others, MS-DIAL 4: accelerating lipidomics using an MS. MS, CCS, and retention time atlas. bioRxiv 2020, 2020.

(7) Broeckling, C. D.; Hoyes, E.; Richardson, K.; Brown, J. M.; Prenni, J. E. Comprehensive Tandem-Mass-Spectrometry Coverage of Complex Samples Enabled by Data-Set-Dependent Acquisition. Anal. Chem. 2018, 90, 8020–8027.

(8) AcquireX Intelligent Data Acquisition Workflow. https://www.thermofisher.com/uk/en/home/industrial/mass-spectrometry/liquid-chromatography-mass-spectrometry-lc-ms/lc-ms-software/lc-ms-data-acquisition-software/acquirex-intelligent-data-acquisition-workflow.html, Accessed: 2020-09-09.

(9) Wandy, J.; Davies, V.; J J van der Hooft, J.; Weidt, S.; Daly, R.; Rogers, S. In silico optimization of mass spectrometry fragmentation strategies in metabolomics. Metabolites 2019, 9, 219.

(10) Pluskal, T.; Castillo, S.; Villar-Briones, A.; Oresic, M. MZmine 2: modular framework for processing, visualizing, and analyzing mass spectrometry-based molecular profile data. BMC Bioinformatics 2010, 11, 395.

(11) Smith, C. A.; Want, E. J.; O’Maille, G.; Abagyan, R.; Siuzdak, G. XCMS: processing mass spectrometry data for metabolite profiling using nonlinear peak alignment, matching, and identification. Anal. Chem. 2006, 78, 779–787.

(12) “Thermo Fisher Scientific”, Thermo Fisher Application Programming Interface. https://github.com/thermofisherlsms/iapi, Accessed: 2020-8-18.

(13) Melnikov, A. D.; Tsentalovich, Y. P.; Yanshole, V. V. Deep Learning for the Precise Peak Detection in High-Resolution LC-MS Data. Anal. Chem. 2020, 92, 588–592.

(14) Tautenhahn, R.; Böttcher, C.; Neumann, S. Highly sensitive feature detection for high resolution LC/MS. BMC Bioinformatics 2008, 9, 504.

(15) Senko, M. W. et al. Novel parallelized quadrupole/linear ion trap/Orbitrap tribrid mass spectrometer improving proteome coverage and peptide identification rates. Anal. Chem. 2013, 85, 11710–11714.

