## Supplementary figures, tables and information for "Rapid Development of Improved Data-dependent Acquisition Strategies"

### Rapid Development of Improved Data-dependent Acquisition Strategies - Supplementary Materials.

#### S1. Peak picking parameters

| Parameters | Settings |
|---|---|
| Mass Detector | Centroid |
| Mass Detector Noise Level | 1000.0 |
| Chromatogram Builder | ADAP |
| Min Group Size | 3 |
| Group Intensity Threshold | 1000.0 |
| Min Highest Intensity | 1000.0 |
| m/z Tolerance (ppm) | 10.0 |
| Deconvolution Method | Wavelets (ADAP) |
| S/N Threshold | 5.0 |
| S/N Estimator | Intensity Window SN |
| Min feature height | 1000.0 |
| Coefficient / Area Threshold | 1.0 |
| Peak Duration Range | (0.0, 5.0) |
| RT Wavelet Range | (0.1, 1.0) |

*Table SI-1: MZMine2 settings used for the peak picking used for evaluation.*

| Parameters | Value |
|---:|:---:|
| algorithm | CentWave |
| ppm | 15 |
| peakwidth | (15, 80) |
| snthresh | 5 |
| noise | 1000 |
| prefilter | (3, 500) |

*Table SI-2: XCMS parameters used in peak picking. All other parameters were XCMS defaults.*

### S2. Scan timing analysis and comparison with vendor method.

Timings were compared for different methods to understand the difference in the number of scans that each method achieves for a given sample on the real MS hardware. Timings for each scan were calculated by looking at the resulting mzML file and computing the difference in scan start times for successive scans. We adopted this approach as we were interested in the full duty cycle, including any processing in ViMMS. Figure SIAA shows these times plotted against the retention time (RT) of the start of the scan for 6 different methods. These methods included 3 methods that worked through the Thermo Fusion IAPI: a full scan controller in C# (csharp_full) bypassing VIMMS and directly using IAPI, a full scan controller in Python using ViMMS (python_full), and a Top-N controller in Python using ViMMS (python_topn). These were compared against 3 vendor methods created using the Xcalibur software suite: a full scan method (fusion_full), a Top-N method (fusion_topn) and a Top-N method with background exclusion (fusion_topn_be). All methods used the same scan parameters, as detailed in the main paper.

The full scan methods in Figure SIAA allow us to compare the comparative speed of the vendor method with those written in C# and Python as part of ViMMS and hence assess any timing overhead introduced by our use of the IAPI. The results show that there is a small difference (around 0.05) seconds between the time taken for a vendor scheduled scan andscans produced by ViMMS. The difference appears to be as a result of the delay in using the IAPI in order to schedule scans. Our implementation requires a bridge between the IAPI (C#) and python (ViMMS). The very small difference between csharp_full and python_full suggest that this additional translation is not costly.

Comparison of the TopN methods is more difficult. The vendor TopN method makes use of some level of parallelisation (Senko *et al.*, 2013). The MS2 scans scheduled from a particular MS1 are started **after** the next MS1 scan has been injected. This means that the time between successive scan start times doesn’t always reflect that actual time a scan takes but, for most MS1 scans, just gives the injection time. This can be seen in the vendor scenario 1 in Figure SI-1.

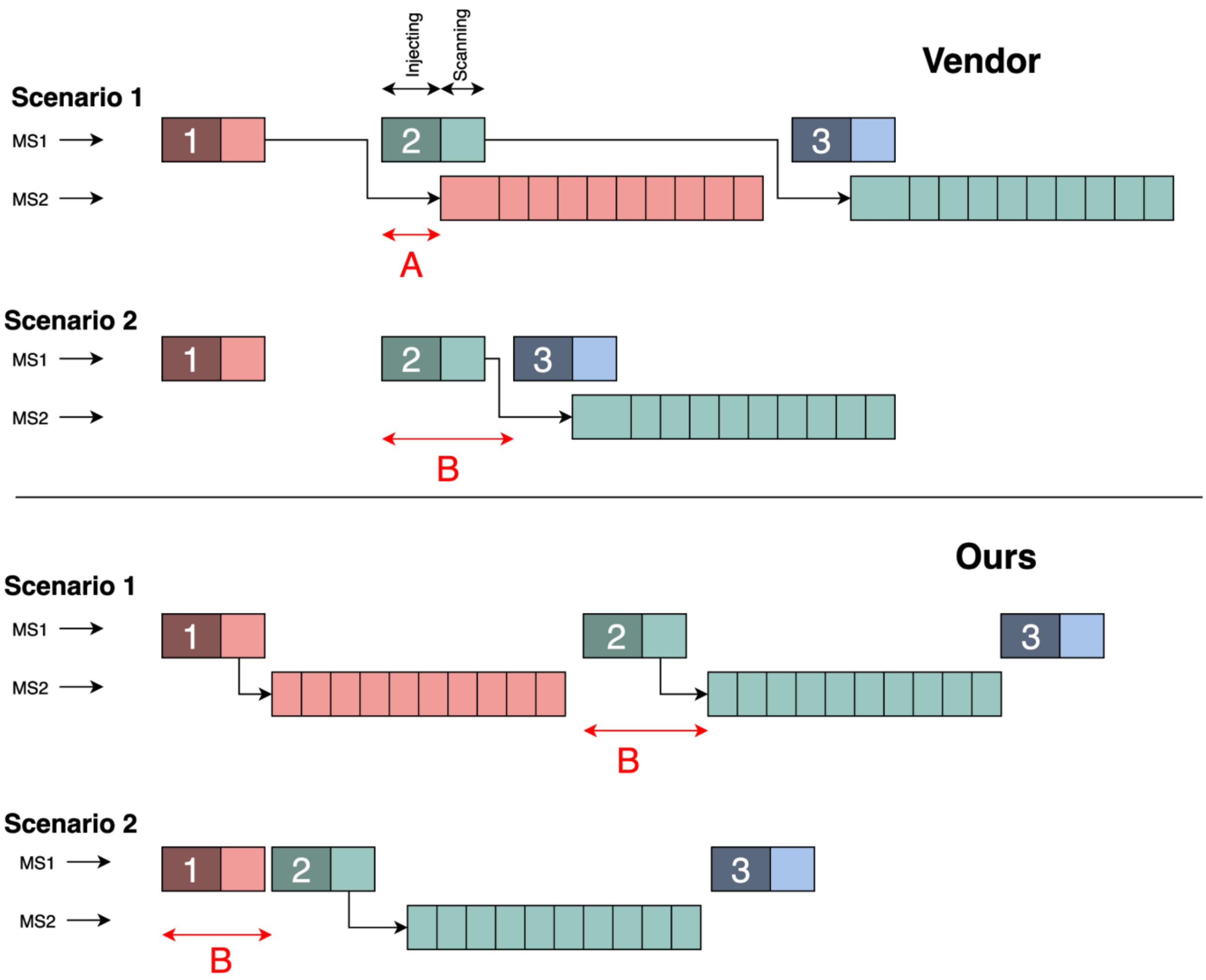


*Figure SI-1: Schematic of scan ordering for vendor (top) and our methods (bottom). In both cases, two scenarios are shown, differing in whether any MS2 scans are scheduled from MS1 scan 1. In Scenario 1 scans are scheduled. In Scenario 2, they are not. Note that this figure is slightly different from that shown in* (Senko et al., 2013) *where they have full parallelisation due to the MS2 scans being performed in the linear ion trap whereas we perform MS2 scans in the orbitrap. This is also why the first MS2 scans in the vendor setup take longer than subsequent ones: they can start injection after the MS1 injection, but scanning cannot start until the MS1 scanning is finished.*

Scenario 1 in the vendor block of Figure SI-1 shows how the first MS2 scan scheduled from MS1 scan 1 starts after the injection of MS1 scan 2 (the time labeled A). Direct comparison of scan times is therefore not possible. However, we can compare the MS1 scan cycle time in situations where no MS2 are scheduled from the previous MS1 scan. This is shown in vendor scenario 2. Here, no scans are scheduled from MS1 scan 1 meaning that the time from the start of MS1 scan 2 to the start of MS1 scan 3 does give us an indication of the MS1 cycle time. As we do not make use of parallelisation in our methods, the times are consistent, regardless of whether or not MS2 scans are scheduled.

Figure SI-2 shows a comparison of MS1 times for TopN methods where vendor times are limited to only the MS1 scan times for scans where no MS2 were scheduled from the previous MS1 scan (time B in vendor scenario 2 in Figure SI-1). We see roughly the same 0.05 second overheard for our methods (compare python_topn with fusion_topn_be). As we only plot vendor instances when no MS2 are scheduled, these are concentrated to the edges of the plot where the chromatogram is less busy. Fusion_topn_be is a machine TopN method with background exclusion turned on. Fusion_topn has no background exclusion and as such we see very few instances when it doesn't fragment and therefore very few points on this plot.

This analysis allows us to conclude that there is very little overhead associated with our use of the IAPI but there may be benefits in the future of implementing similar parallelisation to that provided in the vendor method in our controllers as the parallelisation allows the vendor method to produce many more scans in total. For example, where our TopN controller did 6786 scans in total for QCB (652 MS1 and 6134 MS2), the vendor controller (fusion_topn) with the same scan parameters managed 9901 scans (1299 MS1 and 8602 MS2).

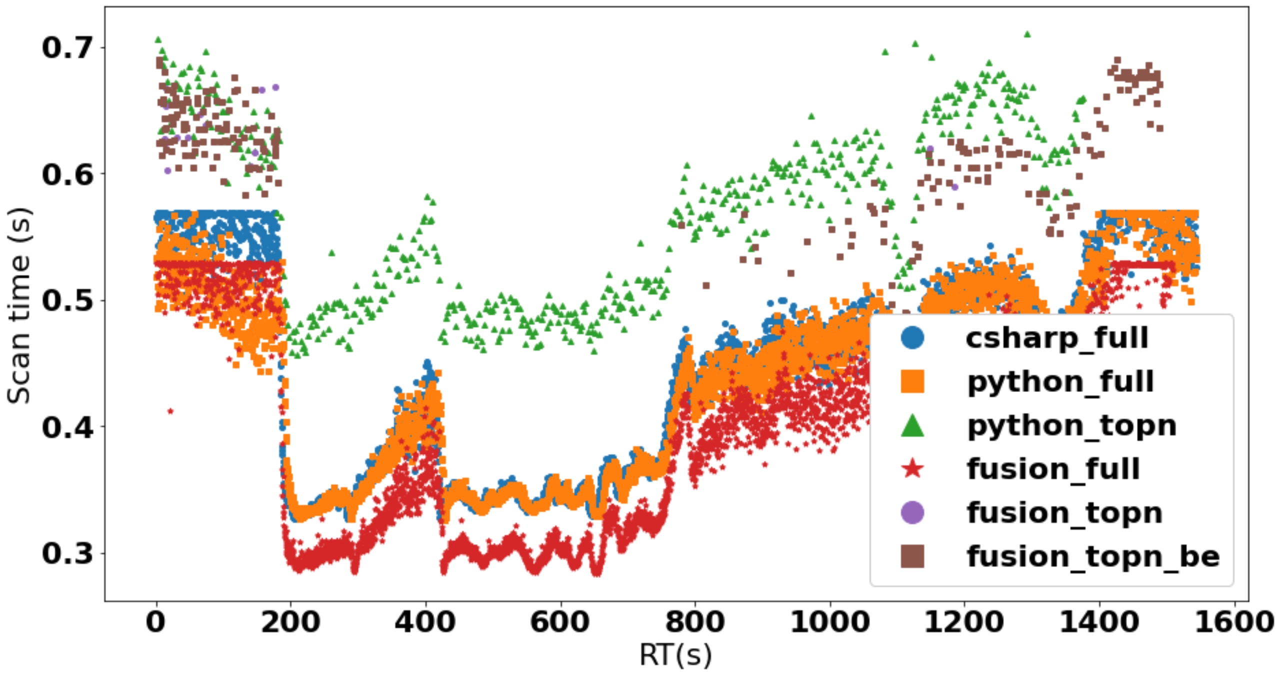


*Figure SI-2: Plot showing the timings for 6 different methods against the retention time (RT) of the scan. For Top-N methods only points that are above the threshold of 0.4 are plotted in order to enhance the visualisation of the results.*

It is interesting to note that this dramatic increase in number of scans does not translate to a large improvement in performance. Table SI-3 shows the coverage of our new controllers as well as the vendor TopN method. We can see that it doesn't outperform any of the proposed methods. It improves over our TopN for QCB but not QCA. This suggests that further development (including some level of parallelisation) could further improve our controllers. Indeed, the optimal results (shown in Table SI-3) using average timings from the vendor method offer an improvement over the optimal results using our TopN timings, particularly for QCB.

| Method | QCB (6267 peaks) | QCA (4481 peaks) |
|---|---|---|
| TopN | 1509 (7) | 1184 (2) |
| WeightedDEW | **2180** (10) | **1534** (2) |
| SmartROI | 1982 (4) | 1281 (-2) |
| SmartROI (shift = 1) | 2047 (4) | 1309 (1) |
| SmartROI (shift = 2) | 2106 (3) | 1318 (1) |
| *Vendor TopN method* | *1919 (5)* | *1158 (-10)* |
| *Optimal (using TopN scan timings)* | *2736* | *1767* |
| *Optimal (using vendor TopN scan timings)* | *3260* | *1795* |

*Table SI-3: method coverage including vendor methods.*

Parellisation must be used with care however. Studying Figure S1, we can see that the parallelisation can lead to a very large time between the survey scan and the later MS2 scans scheduled from it. Consider, for example, the large time from MS1 scan 2 until its MS2 scans. Smaller peaks may have stopped eluting by the time they are needed for fragmentation. Indeed, in our analysis (and other analysis pipelines too), when looking for the precursor m/z for the MS2 scans scheduled from MS1 scan 2, we would actually look to the most recent MS1 (which in this case would be MS1 scan 3). We hypothesis that this may be what is causing the reduction in performance for QCA of the vendor TopN over our TopN, but testing that hypotheses is outside the scope of this work.

### S3. Results for alternative peak picking methods

To ensure our results were not biased by a particular peak picking algorithm, we evaluated performance using two other methods. The Centwave algorithm from XCMS (Smith *et al.*, 2006) and peakonly: a recently proposed method based on a convolutional neural network (Melnikov, Tsentalovich and Yanshole, 2020). Peakonly was run with default parameters whilst XCMS parameters were tuned to give a roughly comparable number of peaks to those found with MzMine. -- the parameters are shown in Table SI-2. Coverage results are shown in Table SI-4. We can see that with both peak picking approaches, our new methods all outperform TopN. For peakonly we see that the SmartROI (with shifts) outperforms the WeightedDEW, but the differences are rather small.

| | XCMS | | Peak Only | |
|---|---|---|---|---|
| **Method** | **QCB (5643 peaks)** | **QCA (3958 peaks)** | **QCB (2510 peaks)** | **QCA (1721 peaks)** |
| TopN | 2012 | 1616 | 1119 | 708 |
| WeightedDEW | **2380** | **1902** | 1375 | 921 |
| SmartROI | 2165 | 1746 | 1360 | 930 |
| SmartROI (shift = 1) | 2238 | 1783 | 1422 | **936** |
| SmartROI (shift = 2) | 2261 | 1705 | **1438** | 920 |

*Table SI-4: Coverage (number of picked peaks fragmented) for each controller for both QCA and QCB, where peaks have been picked using XCMS and PeakOnly. The first five rows show the performance of our (Python) controllers, with the shift set to be the same as Table 1 in the main paper. The final row gives the performance of the instrument method.*

### S4. Computing theoretical fragmentation bounds

We here provide more details on the optimal matching.

A graph `G=(V,E)` is a pair of sets `V` and `E`, where `V` is a set of *vertices* and `E` a set of *edges*. Each edge is an unordered pair of vertices. The vertices that are joined by an edge are known as the *endpoints* of the edge.

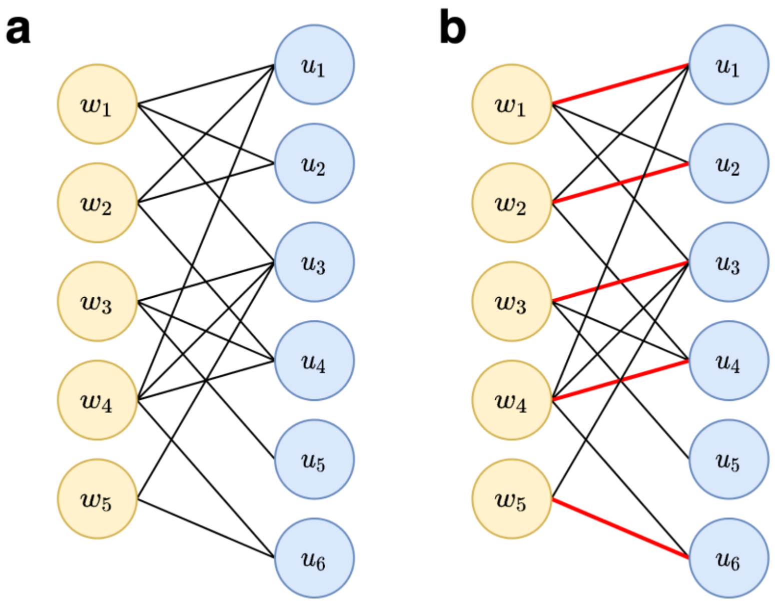


*Figure SI-3: (a) Bipartite graph G and (b) G with a maximum matching shown in red.* $W=\{w_1,w_2,w_3,w_4,w_5\}$ *corresponds to MS2 scans,* $U=\{u_1,u_2,u_3,u_4,u_5,u_6\}$ *corresponds to peak bounding boxes (from peak picking). Edges are drawn for MS2 scans that can be connected to bounding boxes if certain conditions are met to produce the fragmentation events.*

A bipartite graph is a graph for which `V` is the union of two distinct sets, `W` and `U`, say, and for which every edge has one endpoint in `W`, and one endpoint in `V`.  Figure SI-3(a) shows a bipartite graph $G_1$ where $W=\{w_1,w_2,w_3,w_4,w_5\}$ and $U=\{u_1,u_2,u_3,u_4,u_5,u_6\}$. Here `W` corresponds to MS2 scans, while `U` corresponds to the peak boxes. A matching (Erciyes, 2018) of graph $G$ is a subset $M$ of $E$ such that no two edges in $M$ share an endpoint. A maximum matching $M$ is a matching such that no matching $M'$ of $G$ contains more edges. Figure SI-3(b) shows a maximum matching of $G_1$ (red edges). Polynomial time algorithms exist to find maximum matchings. The problem can be solved by adding source and sink nodes and then using an optimised version of the Ford-Fulkerson algorithm (Ford and Fulkerson, 1956) to solve the resulting maximum network flow problem. This algorithm runs in $O(|V|.|E|)$ time. An alternative, faster ($O(\sqrt{|V|}.|E|)$)algorithm is the Hopkropf-Karp algorithm (Hopcroft and Karp, 1973), on which we base our implementation.

### S5. Optimality results

| ***Sample*** | ***Average Scan time (s)*** | | ***Coverage*** | |
|---|---|---|---|---|
| | ***MS1*** | ***MS2*** | ***Observed*** | ***Optimal*** |
| ***QCA (4481 peaks)*** | *0.64* | *0.20* | *1184* | *1767* |
| ***QCB (6267 peaks)*** | *0.59* | *0.20* | *1509* | *2736* |

*Table SI-5: Observed and optimal coverages for QCB and QCA for a TopN method.*

### S6. Scan timings for different controller methods

| | **QCB** | | **QCA** | |
|---|---|---|---|---|
| **Method** | **MS1** | **MS2** | **MS1** | **MS2** |
| Fullscan | 0.43 | NA | 0.48 | NA |
| toTopN | 0.59 | 0.20 | 0.64 | 0.20 |
| SmartROI | 0.70 | 0.20 | 0.71 | 0.20 |
| WeightedDEW | 0.62 | 0.20 | 0.65 | 0.20 |

*Table SI-6: Scan timings for the different controller methods implemented on top of ViMMS. All times are averaged over one injection, in seconds. Time for scan A is computed from the .mzML file as the difference between the scan start time of scans A and A+1 and therefore include both the time required to scan and process. As expected, MS2 times are constant, but MS1 times vary depending on the computation required to process the MS1 scan. Fullscan requires no processing and is therefore the quickest. SmartROI requires the most computation and is, as expected, the slowest.*

### S7. *In-silico* optimisation of controllers

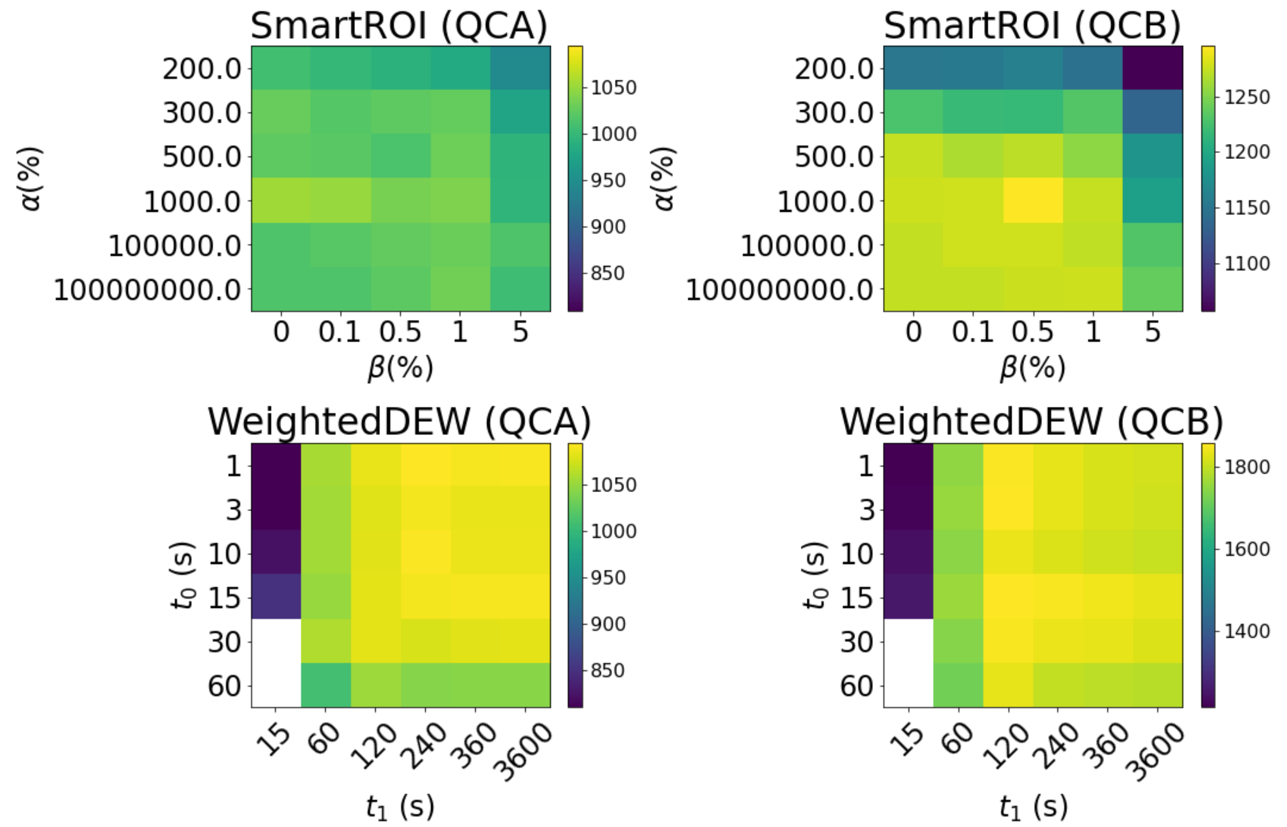


*Figure SI-4: Heatmaps of simulated coverage for SmartROI (top) and WeightedDEW (bottom). The simulated results could be used to suggest appropriate parameters to use for validation on the actual instruments. White cells in the WeightedDEW results indicate invalid parameter combinations.*

### References


Erciyes, K. (2018) 'Matching', in Erciyes, K. (ed.) *Guide to Graph Algorithms: Sequential, Parallel and Distributed*. Cham: Springer International Publishing, pp. 263–303. doi: 10.1007/978-3-319-73235-0_9.

Ford, L. R. and Fulkerson, D. R. (1956) 'Maximal Flow Through a Network', *Canadian Journal of Mathematics. Journal Canadien de Mathematiques*. Cambridge University Press, 8, pp. 399–404. doi: 10.4153/CJM-1956-045-5.

Hopcroft, J. E. and Karp, R. M. (1973) 'An $n^{5/2} $ Algorithm for Maximum Matchings in Bipartite Graphs', *SIAM Journal on Computing*. Society for Industrial and Applied Mathematics, 2(4), pp. 225–231. doi: 10.1137/0202019.

Melnikov, A. D., Tsentalovich, Y. P. and Yanshole, V. V. (2020) 'Deep Learning for the Precise Peak Detection in High-Resolution LC-MS Data', *Analytical chemistry*, 92(1), pp. 588–592. doi: 10.1021/acs.analchem.9b04811.

Senko, M. W. *et al.* (2013) 'Novel parallelized quadrupole/linear ion trap/Orbitrap tribrid mass spectrometer improving proteome coverage and peptide identification rates', *Analytical chemistry*, 85(24), pp. 11710–11714. doi: 10.1021/ac403115c.

Smith, C. A. *et al.* (2006) ‘XCMS: processing mass spectrometry data for metabolite profiling using nonlinear peak alignment, matching, and identification’, *Analytical chemistry*, 78(3), pp. 779–787. doi: 10.1021/ac051437y.
